# Chronic unpredictable stress promotes cell-specific plasticity in prefrontal cortex D1 and D2 pyramidal neurons

**DOI:** 10.1101/565499

**Authors:** EM Anderson, D Gomez, A Caccamise, D McPhail, M Hearing

**Affiliations:** Department of Biomedical Sciences, Marquette University, Milwaukee, WI, 53233

**Keywords:** prefrontal cortex, stress, intrinsic excitability, miniature postsynaptic currents, affective behavior, dopamine receptor

## Abstract

Exposure to unpredictable environmental stress is widely recognized as a major determinant for risk and severity in neuropsychiatric disorders such as major depressive disorder, anxiety, schizophrenia, and PTSD. The ability of ostensibly unrelated disorders to give rise to seemingly similar psychiatric phenotypes highlights a need to identify circuit-level concepts that could unify diverse factors under a common pathophysiology. Although difficult to disentangle a causative effect of stress from other factors on medial prefrontal cortex (PFC) dysfunction, a wealth of data from humans and rodents demonstrates that the PFC is a key target of stress. The present study sought to identify a model of chronic unpredictable stress (CUS) which induces affective behaviors in C57BL6J mice and once established, measure spike firing and the ability to evoke an action potential in mPFC layer 5/6 pyramidal neurons. Adult male mice received 2 weeks of ‘less intense’ stress or 2 or 4 weeks of ‘more intense’ CUS followed by sucrose preference for assessment of anhedonia, elevated plus maze for assessment of anxiety and forced swim test for assessment of depressive-like behaviors. Our findings indicate that more intense CUS exposure results in increased anhedonia, anxiety, and depressive behaviors, while the less intense stress results in no measured behavioral phenotypes. Once a behavioral model was established, mice were euthanized approximately 21 days post-stress for whole-cell patch clamp recordings from layer 5/6 pyramidal neurons in the prelimbic (PrL) and infralimbic (IL) cortices. No significant differences were initially observed in intrinsic cell excitability in either region. However, post-hoc analysis and subsequent confirmation using transgenic mice expressing tdtomato or eGFP under control of dopamine D1- or D2-type receptor showed that D1-expressing pyramidal neurons (D1-PYR) in the PrL exhibit reduced thresholds to fire an action potential (increased excitability) but impaired firing capacity at more depolarized potentials, whereas D2-expressing pyramidal neurons showed an overall reduction in excitability and spike firing frequency. Examination of synaptic transmission showed that D1- and D2-PYR in exhibit differences in basal excitatory and inhibitory signaling under naïve conditions. In CUS mice, D1-PYR showed increased frequency of both miniature excitatory and inhibitory postsynaptic currents, whereas D2-PYR only showed a reduction in excitatory currents. These findings demonstrate that the intrinsic physiology and synaptic regulation of D1- and D2-PYR subpopulations differentially undergo stress-induced plasticity that may have functional implications for stress-related pathology, and that these adaptations may reflect unique differences in basal properties regulating output of these cells.

## 1.0 Introduction

Exposure to unpredictable environmental stress is widely recognized as a major determinant of risk and severity in neuropsychiatric disorders such as major depressive disorder (MDD), anxiety, and post-traumatic stress disorder (Bale, 2005; Kendler, Karkowski, & Prescott, 1998, 1999; Moghaddam & Javitt, 2012). The medial prefrontal cortex (mPFC) is intricately involved in cognitive performance, as well as top-down regulation of affect and stress responsivity (Anisman & Matheson, 2005; Clark, Chamberlain, & Sahakian, 2009; Fossati, Amar, Raoux, Ergis, & Allilaire, 1999; Herman, Ostrander, Mueller, & Figueiredo, 2005; Keedwell, Andrew, Williams, Brammer, & Phillips, 2005; Krishnan & Nestler, 2008; Miller & Lewis, 1977; Murphy et al., 1999; Murrough, Iacoviello, Neumeister, Charney, & Iosifescu, 2011; Radley, Arias, & Sawchenko, 2006; Sullivan, 2004; Treadway & Zald, 2011).

Functional integrity of mPFC information processing and downstream communication relies on a dynamic balance of excitatory and inhibitory signaling, with disruptions in this balance implicated in stress-related pathologies including flattened affect (anhedonia), anxiety-like behavior, and impaired cognition (Gandal et al., 2012; Holmes & Wellman, 2009; Matsuo et al., 2007; Sohal, Zhang, Yizhar, & Deisseroth, 2009; Yizhar et al., 2011). Structural modifications in pyramidal neurons (PYR) – the principle output neurons in the mPFC – have long been thought to play a critical role in stress-induced cortical dysfunction, however to date only a handful of studies have examined the impact of this reorganization on neurotransmission and cellular physiology, with even fewer examining the cell-specific locus of these adaptations (McEwen & Morrison, 2013; McKlveen et al., 2016; Radley et al., 2005; Radley, Rocher, et al., 2006; Shansky & Morrison, 2009; Urban & Valentino, 2017).

Growing evidence indicates that distinctions in molecular (e.g., ion channels, receptors), neurophysiology, and anatomical connectivity endow specific subpopulations of PYR with unique properties to integrate input and communicate information downstream (Brown & Hestrin, 2009; Degenetais, Thierry, Glowinski, & Gioanni, 2002; Dembrow, Chitwood, & Johnston, 2010; Gee et al., 2012; Kim, Ahrlund-Richter, Wang, Deisseroth, & Carlen, 2016; Seong & Carter, 2012; Sohal et al., 2009; Yang, Seamans, & Gorelova, 1996). For example, recent evidence indicates that PYR neurons expressing either the dopamine D1 (D1-PYR) or D2 (D2-PYR) receptor exhibit distinct neurophysiological properties and synaptic innervation, and in some instances subcortical projection targets, that likely define how they contribute to behavior and undergo experience-induced plasticity (e.g., stress) (Anastasiades, Boada, & Carter, 2018; Benes, Vincent, & Molloy, 1993; Gee et al., 2012; Santana, Mengod, & Artigas, 2009; Seong & Carter, 2012; Xu & Yao, 2010). As these cortical networks likely provide a neuroanatomical framework for complex regulation of behavior (Brumback et al., 2018; Gaspar, Bloch, & Le Moine, 1995; Gee et al., 2012; Jenni, Larkin, & Floresco, 2017; Santana et al., 2009; Seong & Carter, 2012; Vincent, Khan, & Benes, 1993), a critical step towards understanding how stress influences behavior include identifying the selectivity of stress-induced plasticity and associated mechanisms (Jenni et al., 2017).

The current study set out to establish a model of chronic unpredictable stress (CUS) in C57BL6J mice that induces consistent affective behaviors as well as determine how CUS differentially impacts mPFC D1- and D2-expressing PYR neuron intrinsic physiology and plasticity. Findings from this study have implications to increase understanding on the influence that chronic stress exposure has on enduring changes in mPFC that may contribute to alterations in behaviors, including behaviors within anxiety disorders and major depression.

## 2. Materials and Methods

### 2.1 Animals

Adult male mice (PD51-74) were a combination of wild-type (C57BL/6J) bred in house or purchased from Jackson Laboratories, heterozygous bacterial artificial chromosome (BAC) transgenic mice (Jackson Laboratories) expressing tdtomato or eGFP expression, or double transgenics expressing tdtomato and eGFP driven by either DR1 (drd1a-tdtomato) or DR2 (drd2-eGFP) dopamine receptors. Recordings performed from single transgenics expressing only tdtomato driven by DR1 were used as the tdtomato signaling is greater in cortical neurons compared to eGFP and also exhibits decreased photobleaching compared to eGFP, therefore cells were identified as D1+ or D1-. Mice were maintained in a temperature and humidity-controlled room with all procedures approved by the Institutional Animal Care and Use Committee at Marquette University.

### 2.2 Chronic Unpredictable Stress

Mice were exposed to two weeks of less intense (LI) stress or exposed to two or four weeks of more intense (MI) stress (Figure 1). To increase stress intensity, the level of unpredictability was increased by further varying the times, durations, and locations as well as combining stressors (e.g. cage tilt in cold room) and using MI stressors with increased frequency.

**Figure 1.**
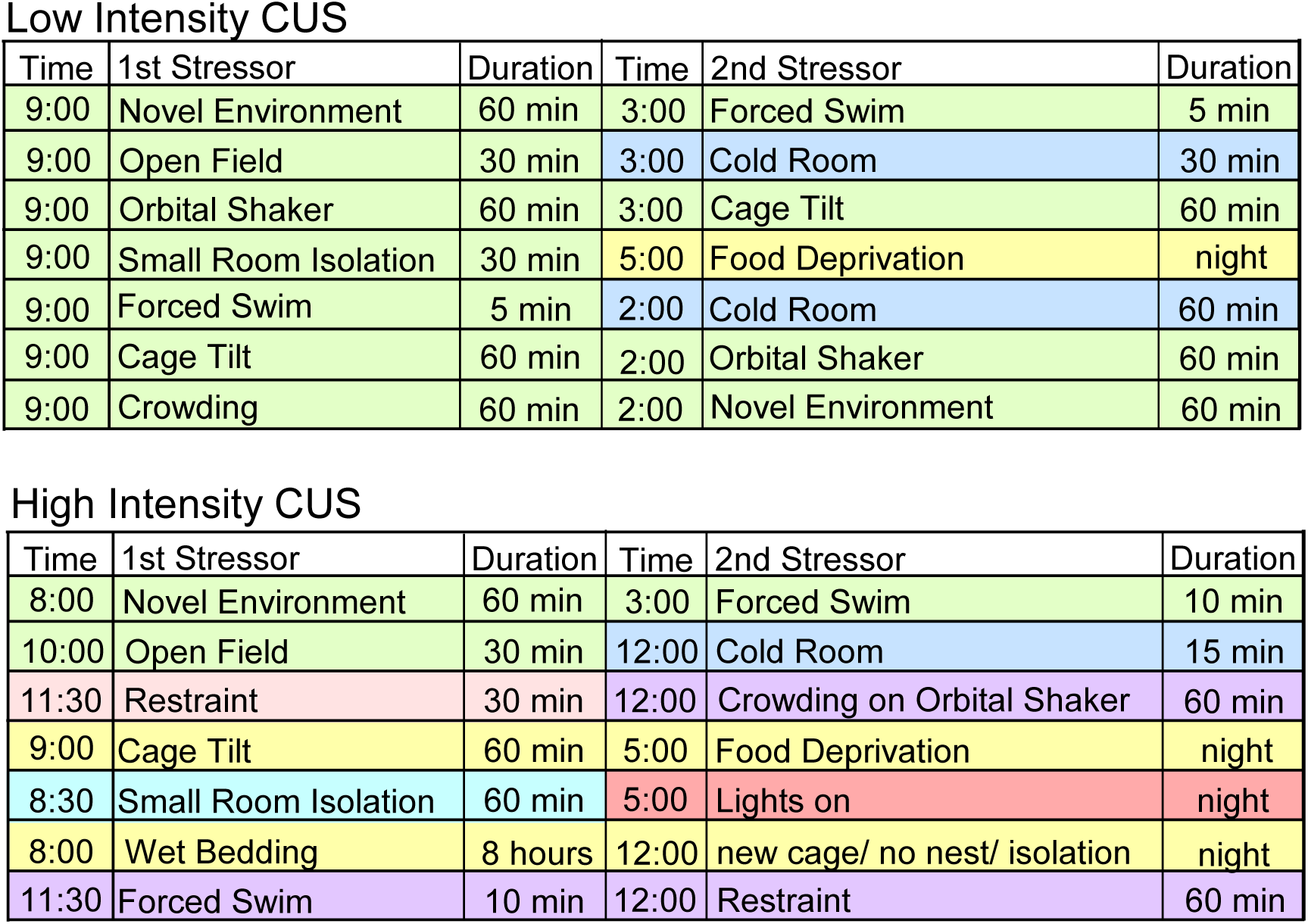
One week sample of unpredictable stressors of various durations, intensities, and in various locations (green= stress room A, red= stress room B, purple= stress room C, blue= cold room, yellow= colony). Mice received two weeks of less intense stress (top) or two or four weeks of more intense stress (bottom).

### 2.3 Behavioral Testing

#### Sucrose preference

In a subset of mice, sucrose preference was assessed as a measure of anhedonia (Forbes, Stewart, Matthews, & Reid, 1996; Willner, Towell, Sampson, Sophokleous, & Muscat, 1987). The evening of the last stress exposure (Figure 2A), mice were provided two separate bottles that were weighed, one containing 1% sucrose solution and the other containing tap water. The mouse had *ad libitum* access to food and both bottles overnight. The following morning, the bottles were removed and reweighed. Percent sucrose consumed was calculated as the amount of sucrose water consumed divided by the amount of sucrose water consumed plus the amount of tap water consumed.

**Figure 2.**
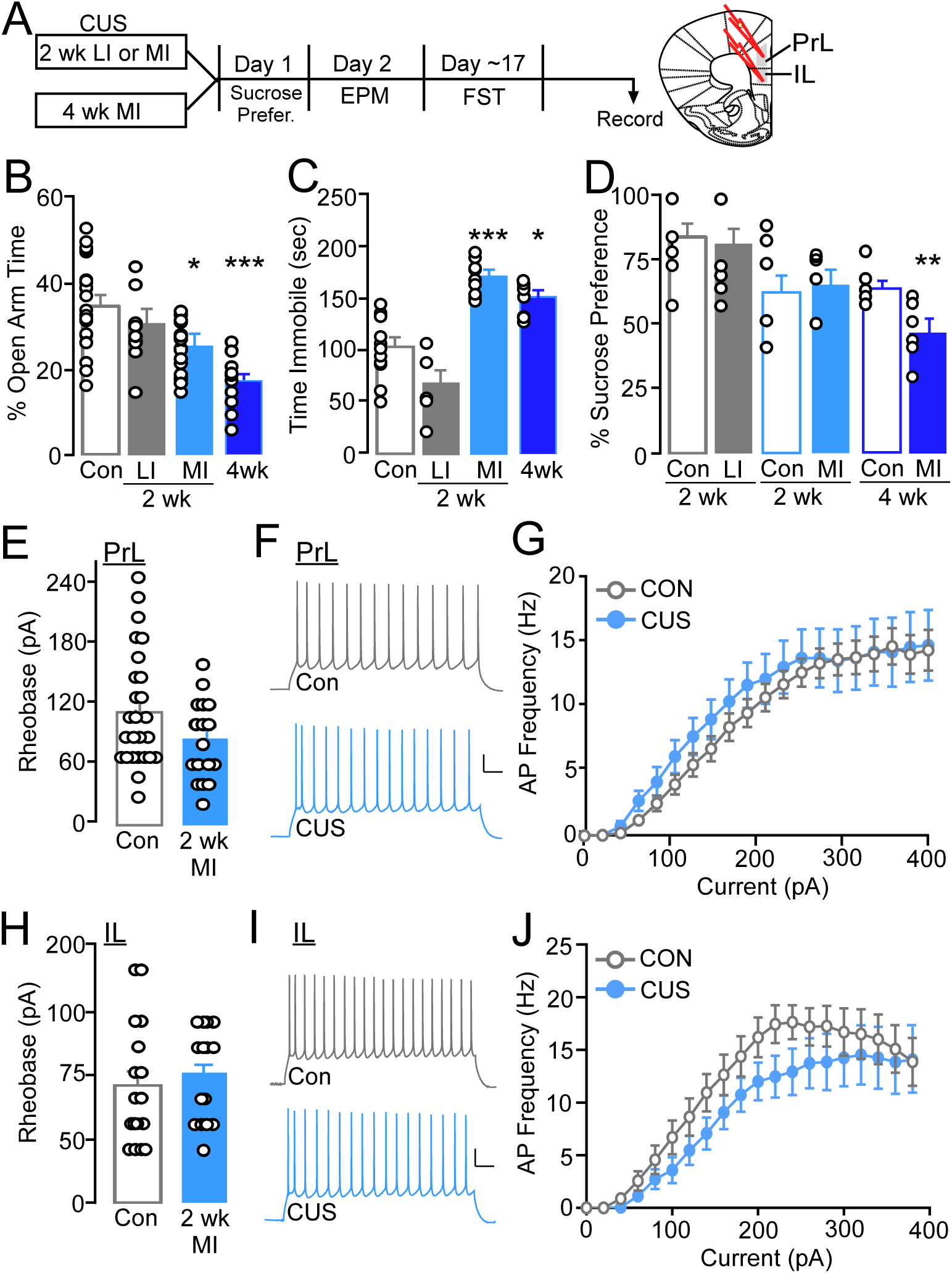
(A) Mice received either two weeks of less or more intense stress or four weeks of more intense stress following by behavioral testing and slice electrophysiology in the PrL or IL. (B) Percent time in the open arm of the elevated plus maze. Mice exposed to two or four weeks of more intense stress had significant reductions in percent open arm time (*N*= 9-29/group). (C) Mice exposed to two or four weeks of more intense stress had significant increases in time immobile during the forced swim test (*N*= 6-14/group). (D) Percent sucrose consumed during an overnight preference test, only mice exposed to four weeks of more intense stress had significant decrease in preference (*N*=6-18/group). (E) There were no differences in the current required to evoke an action potential (rheobase) in L5/6 PrL PYR neurons from control and mice exposed to two week more intense stress (*n*=17-23/group, *N*=9-11/group). (F) Spiking elicited during a one second 260pA current injection in L5/6 PrL PYR from control (top) and 2 week MI CUS (bottom) mice. (G) There were no differences in the current-spike plots for control and CUS L5/6 PrL PYR neurons. (H) No differences in rheobase in L5/6 IL PYR neurons from control and 2 week MI CUS mice (*n*=16-17/group, *N*=8-9/group). (I) Spiking elicited during a one second 260pA current injection in L5/6 IL PYR from control (top) and 2 week MI CUS (bottom) mice. (J) No differences in current-spike plots for control and 2 week MI CUS L5/6 IL PYR neurons. (scale bar, 20pA/500msec). *p≤0.05 versus CON, ***p<0.001 versus CON.

#### Elevated plus maze

On the day following the last stress exposure (Figure 2A), a subset of mice were tested for anxiety-like behaviors using an elevated plus maze (EPM; San Diego Instruments). The EPM consisted of two opposite open arms (H: 15.25” W: 2.0” L: 26.0”) with lights (∼50 lux) and camera mounted above to monitor and record behavior. Individual trials lasted 5 minutes beginning with the mouse being placed in the center of the maze facing an open arm. Following testing, the maze was cleaned with 70% ethanol and allowed to dry completely between each trial. Behavior was recorded using AnyMaze (Stoelting Co.) tracking software. Percent time in the open arms was calculated as total time in the open arms divided by total time in the maze.

#### Forced swim test

The forced swim test (FST) can be used to assess depression-like behavior or active (i.e. escape behavior) versus passive (i.e. immobility) coping strategies. In the current study, the apparatus was a transparent glass cylinder (7” diameter). Cylinders were filled with 25 ± 2°C water to a 10-15 cm depth to prevent the mouse from touching the bottom. Each mouse was individually habituated for two minutes, with behavioral monitoring occurring during a subsequent four-minute test during which the time immobile (sensitivity: 85%, minimum immobility period: 250 ms) using a side mounted Firewire camera directly facing the cylinder and AnyMaze tracking software. Following testing, the mouse was immediately dried and kept in a warming holding cage.

### 2.4 Slice electrophysiology

Acute slice electrophysiology was performed 20-26 days after the final stress exposure (Figure 2A and 3A). Mice were anesthetized with an overdose of isoflurane (Henry Schein), decapitated, and the brain removed and put in ice-cold solution containing 229mM sucrose, 1.9mM KCl, 1.2mM NaH_2_PO_4_, 33mM NaHCO_3_, 10mM glucose, 0.4mM ascorbic acid, 6mM MgCl_2_, and 0.5mM CaCl_2_ oxygenated using 95% O_2_ 5% CO_2_. Coronal slices (300µm) containing the mPFC were sliced in the ice-cold sucrose solution using a vibratome (Leica VT1000S) and then incubated at 31°C for ten minutes in a solution containing 119mM NaCl, 2.5mM KCl, 1mM NaH_2_PO_4_, 26.2mM NaHCO_3_, 11mM glucose, 0.4mM ascorbic acid, 4mM MgCl_2_, and 1mM CaCl_2_ and further incubated a minimum of 35 minutes at room temperature.

**Figure 3.**
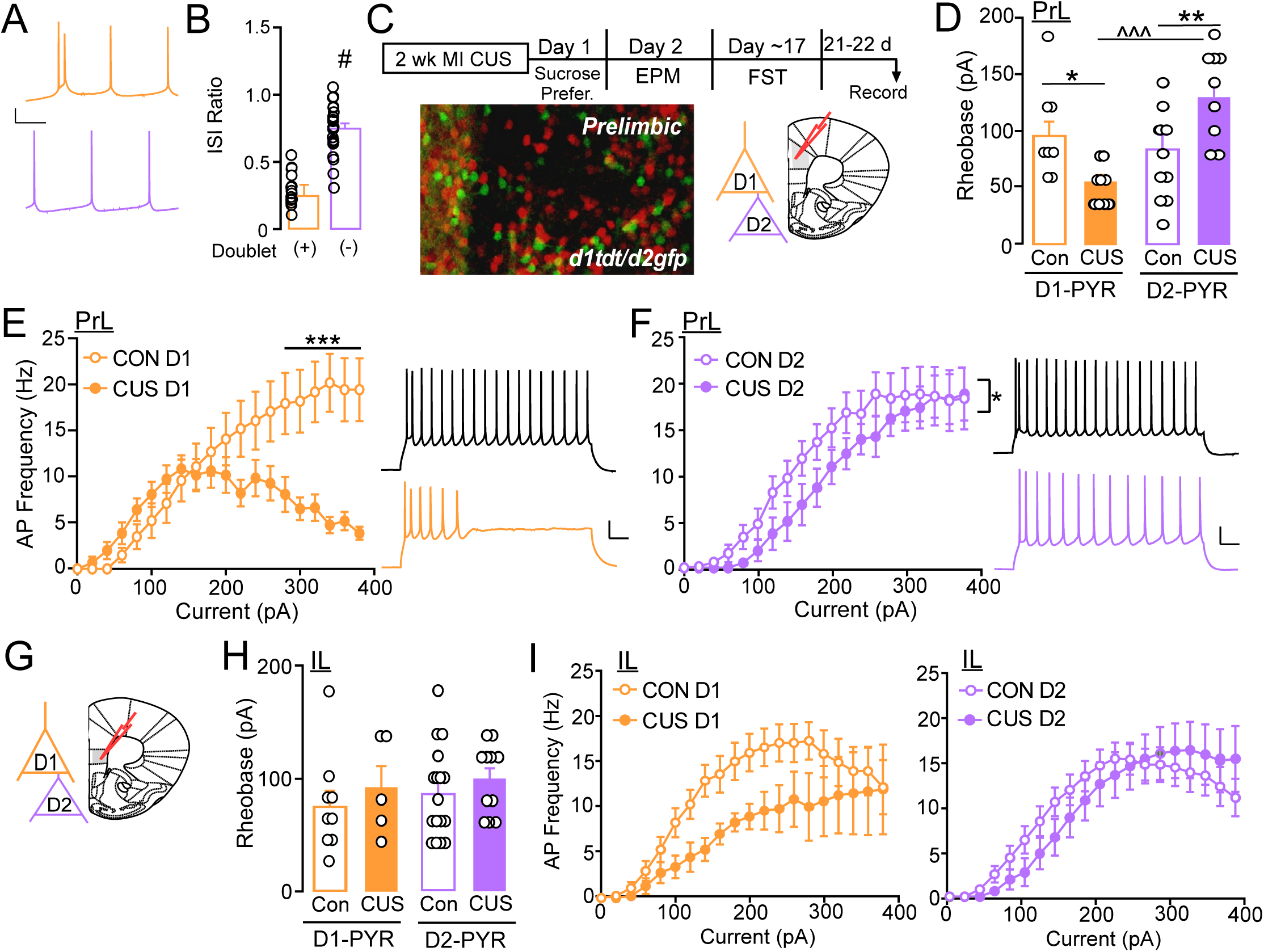
(A) Putative D1-PYR were characterized by a spike ‘doublet’ (orange; top) whereas D2-PYR were characterized by lack of the doublet (purple; bottom). (B) The presence of a spike doublet was positively correlated with the interspike interval (ISI) ratio. (C) Mice received no stress or two weeks of more intense stress with a portion of mice receiving behavioral testing. Image of D1 (red) and D2 (green) fluorescent cells in PrL of double heterozygous BAC transgenic mice (image was modified and enhanced for contrast). (D) Mean current required to evoke an action potential in L5/6 D1-PYR PrL neurons was significantly lower from CUS mice compared to control mice while D2-PYR required significantly more current in CUS mice compared to control. (*n*=8-11/group, *N*=4-6/group). (E) Mean current-spike plots for control and CUS L5/6 PrL D1-PYR neurons show lower firing at more depolarized potentials in CUS mice. (F) CUS L5/6 PrL D2-PYR neurons had overall lower spike firing during current injections compared to D2-PYR neurons from control mice. (G) Representative coronal section showing IL region of recordings. (H) Mean current required to evoke an action potential in L5/6 D1 and D2-PYR neurons in the IL was similar for control and CUS mice. (I) There were no differences in current-spike plots for control and CUS L5/6 IL D1- or D2-PYR neurons. #p<0.001 versus presence of doublet, *p≤0.05, **p<0.01. ***p<0.001 versus CON vs CUS; ^^^p<0.001 CUS vs CUS.

During whole-cell recordings, slices were continuously perfused with oxygenated aCSF (125mM NaCl, 2.5mM KCl, 25mM NaHCO_3_, 10mM glucose, 0.4mM ascorbic acid, 1.3mM MgCl_2_, 2mM CaCl_2_) at a temperature of 29°C-33°C using a gravity-fed perfusion system with a flow rate of ∼2 ml/min. All recordings were performed with adequate whole-cell access (Ra<40 MΩ). Data was filtered at 2kHz and sampled at 5kHz for current-clamp recordings and 20kHz for miniature postsynaptic current recordings using a Sutter Integrated Patch Amplifier (IPA) with Igor Pro (Wave Metrics, Inc.) data acquisition software.

Deep layer 5/6 PYR neurons were identified based on morphology and/or the presence of fluorescence, as well as physiologically by capacitance (PrL >100pf; IL >75pF) and minimum resting membrane potential (−55mV). For rheobase and action potential firing, borosilicate glass pipettes were filled with internal solution containing140mM K-Gluconate, 5.0mM HEPES, 1.1mM EGTA, 2.0mM MgCl_2_, 2.0mM Na_2_-ATP, 0.3mM Na-GTP, and 5.0mM phosphocreatine (pH 7.3, 290mOsm). Miniature excitatory (mEPSCs) and inhibitory (mIPSCs) postsynaptic currents were recorded using borosilicate glass pipettes (Sutter Instruments; 2.5-4.5 MΩ) filled with a cesium-based internal solution (120mM CsMeSO4, 15mM CsCl, 10mM TEA-Cl, 8mM NaCl, 10mM HEPES, 5mM EGTA, 0.1mM spermine, 5mM QX-314, 4mM ATP-Mg, and 0.3mM GTP-Na). mEPSCs and mIPSCs were recorded in the presence of 0.7mM lidocaine to block Na+-dependent at −72 and 0 mV, respectively.

#### Data analysis

Statistical significance was determined using independent-samples t-test, analysis of variance (ANOVA), two-way ANOVA, or two-way RM ANOVA where appropriate/indicated. Bonferroni post-hoc comparisons were conducted when necessary. Data points +/-2 standard deviations from the mean were excluded which included a total of two control and one stress mouse from EPM analysis. A total of three cells from assessment of unidentified PrL PYR and one D2 putative cell recording were excluded, but none were from mice excluded based on behavior. Data was analyzed using SPSS 24 (IBM Statistics) or SigmaPlot, and graphed using GraphPad Prism. Experimental sample size is presented as *n* for the number of cells and *N* for the number of mice.

## 3. Results

### 3.1 Influence of CUS intensity and duration on affective behavior

The influence of chronic stress exposure on affective behaviors related to anxiety, depression, and anhedonia have been well established in *rats*, however prior research has indicated that the most widely used mouse strain (i.e. C57/BL6) exhibit attenuated stress-induced neuroendocrine responsivity and behavioral deficits compared to other strains (e.g. Balb/c; DBA/2J) (Anisman, Hayley, Kelly, Borowski, & Merali, 2001; Anisman, Lacosta, Kent, McIntyre, & Merali, 1998; Razzoli, Carboni, Andreoli, Ballottari, & Arban, 2011; Razzoli, Carboni, Andreoli, Michielin, et al., 2011; Savignac et al., 2011). Although recent work established a chronic unpredictable stress (CUS) protocol in mice that results in behavioral phenotypes, these protocols required either 4 or 8 weeks of exposure (Monteiro et al., 2015). In an attempt to identify a more efficient protocol that will increase throughput in mice and produce reliable deficits in commonly examined affect-related behavior, initial studies examined three CUS protocols that varied in intensity/predictability as well as duration.

#### 3.1.1 Elevated Plus Maze

To identify effects of variable stress intensity on anxiety-like behavior, mice underwent testing in an elevated plus maze (EPM). There were no significant differences between the three control groups and they were therefore combined (F_(2, 26)_= 0.67, p=0.52). There were significant differences comparing the four conditions (F_(3, 68)_= 11.53, p<0.001), an effect that was not due to differences in locomotion as assessed by combining the number of open, closed, and center arm entries (F_(3, 68)_= 2.52, p=0.07). Bonferroni post-hoc comparisons indicate that less intense stress was similar to control, but both two weeks and four weeks of MI stress had significantly less percent open arm time compared to non-stressed controls [*CON:* 34.12 ± 2.04%; *LI:* 28.60 ± 2.85%, p=0.67; *two week:* 25.95 ± 1.78%, p=0.02; *four week:* 17.16 ± 1.74%, p<0.001; Figure 2B].

#### 3.1.2 Forced Swim

Mice were tested in a forced swim test to determine if chronic stress intensity alters depression-like behavior - a measure previously shown to respond to anti-depressant treatment (Castagne, Moser, Roux, & Porsolt, 2010; Kara, Stukalin, & Einat, 2018) or induces immobility, a passive coping strategy (Commons, Cholanians, Babb, & Ehlinger, 2017; Molendijk & de Kloet, 2015). No differences were observed between control groups, thus they were combined (F_(2, 14)_= 2.58, p=0.12). Similar to measures of anxiety-like behavior, there was a significant difference comparing the four conditions (F_(3, 41)_= 17.81, p<0.001). Exposure to two weeks of LI stress did not alter time spent immobile during the forced swim test however both lengths of the MI stressors significantly increased immobility time [*CON:* 113.16 ± 10.56s, *LI*: 75.38 ± 17.96s, p=0.16; *two week*: 181.30 ± 6.43s, p<0.001; f*our week*: 154.56 ± 9.99s, p=0.05; Figure 2C).

#### 3.1.3 Sucrose Preference

To assess for the potential influence of chronic stress exposure on anhedonia, the percent of sucrose water to tap water consumed over a 24 h period was compared (Figure 2D). There were significant differences in sucrose consumption across the three control groups and therefore were not combined (F_(2,24)_= 6.38, p<0.01). Mice that were exposed to two weeks of chronic stress whether it was LI or MI show similar preference for sucrose compared to control mice [*LI CON:* 83.57 ± 4.16%; *LI:* 81.48 ± 5.23%, t_(18)_= 0.31, p=0.76; *two week CON:* 62.54 ± 6.00%, *two week MI:* 65.28 ± 5.01%, t_(13)_= −0.32, p=0.75]. Conversely, mice that were exposed to four weeks of MI CUS showed a significant reduction in the percent of sucrose compared to water that was consumed, indicating increased anhedonia [*CON:* 63.40 ± 3.08%, *four week:* 46.43 ± 4.78%, t_(10)_= 2.99, p=0.01].

### 3.2 Region specific effects of CUS on randomly selected L5/6 mPFC pyramidal neurons

Subregions of the mPFC demonstrate distinct patterns of connectivity and make dissociable contributions to behavior, including those related to affect (Dalley, Cardinal, & Robbins, 2004; Marquis, Killcross, & Haddon, 2007; Vertes, 2004). Previous findings have shown that intrinsic properties (e.g., excitability) are altered during an acute post-stress period (24 h) following repeated resident-intruder social stress in *mid-adolescent*, but not adult male mice (Urban & Valentino, 2017), however it is unclear whether a CUS model of exposure alters PYR physiology and/or if these effects persist in adult males. Using whole-cell current clamp recordings, we assessed the threshold of current needed to reach depolarization threshold to fire an initial action potential (rheobase) in L5/6 PYR of the PrL and IL regions of the mPFC 20-26 days post stress. The pattern (frequency) of action potential firing in response to increasing current amplitude injections was also assessed to determine whether intrinsic firing properties of these neurons was altered following 2 week MI CUS (Figure 2A). As LI stress did not alter measures of affective behavior, and increased intensity for both lengths of exposure prompted similar deficits – particularly performance in the FST that aligned temporally with recordings – electrophysiology measures were focused on mice undergoing two weeks of increased CUS intensity for all subsequent studies.

Examination of rheobase showed no significant difference between unidentified PrL neurons in stress naïve and 2 week MI CUS exposed mice [t_(42)_= 1.26, p=0.22; CON: 102.22 ± 10.03 pA, CUS: 83.53 ± 9.74pA; Figure 2F]. Examination of current-spike relationship curves showed that CUS did not significantly alter the number of action potentials produced by increasing (20 pA) current steps (two-way repeated-measures ANOVA: condition (control, CUS) (F_(1, 42)_= 0.42, p=0.52); interaction: F_(19, 798)_= 0.45, p=0.98; Figure 2E and 2G). Similarly, L5/6 PYR in the IL did not show effects of 2 week MI CUS on rheobase [t_(29)_= −0.48, p=0.64; CON: 91.25 ± 12.11pA, CUS: 98.67 ± 9.45pA; Figure 2I] or action potential firing frequency (condition (F_(1, 29)_= 2.174, p=0.15); interaction (F_(19, 551)_= 0.55, p=0.94; Figure 2H and 2J). Taken together, these findings indicate that 2 week MI CUS does not produce a *global* effect on PrL or IL PYR intrinsic excitability or that these adaptations do not persist three weeks following conclusion of stress.

### 3.3 Effects of CUS on mPFC D1- and D2-expressing pyramidal neuron intrinsic excitability

PYR express either the dopamine D1-(D1-PYR) or D2 (D2-PYR) receptor, with little overlap (Gaspar et al., 1995; Gee et al., 2012; Santana et al., 2009; Vincent et al., 1993), and may define how they undergo experience-induced plasticity and/or their contribution to behavior (Gee et al., 2012; Jenni et al., 2017; Santana et al., 2009; Seong & Carter, 2012). To determine whether the lack of effect on excitability in randomly selected PYR following stress reflected cell-specific adaptations, we initially reanalyzed rheobase and spike-firing data in PrL PYR based on previously identified physiological characteristics shown to align with D1- and D2-PYR populations (Lee et al., 2014; Seong & Carter, 2012). Briefly, neurons were classified by the presence of a spike “doublet” (putative D1+) or not (putative D1-; Figure 3A). In agreement with previous work, the presence of a doublet was positively correlated (r=0.76, p<0.001) with the inter-spike interval (ISI) ratio of the first and second action potential and the fourth and fifth action potential in a train of at least five action potentials during current-step injections ((AP_2_-AP_1_)/(AP_5_-AP_4_) = ISI Ratio; Figure 3B). In stress naïve mice, rheobase values of PrL putative D1-PYR did not differ compared to values observed in putative D2-PYR (D1-PYR: 93.33 ± 13.78 pA; D2-PYR: 110.00 ± 15.51 pA; t_(24)_= −0.79, p=0.44; data not shown), indicating that baseline excitability of PYR is not defined by the presence of D1- or D2-receptors. Conversely, putative D1-PYR neurons from stress-exposed mice, albeit statistically underpowered, exhibited lower threshold to fire an action potential compared to putative D2-PYR (D1-PYR: 53.33 ± 6.67 pA; D2-PYR: 90.00 ± 11.04 pA; t_(13)_= −2.84, p=0.01; data not shown).

To confirm our initial findings, we used bacterial artificial chromosome (BAC) transgenic mice expressing tdTomato and/or enhanced green fluorescent protein (eGFP) in D1R- and D2R- PYR, respectively (Figure 3C). These mice were also ran through behavioral assessments and did not differ compared to wild-type mice therefore combined and presented above. A main effect of treatment but not cell-type on resting membrane potential (RMP) (Treatment: F_(1, 34)_=4.303, p=0.046). *Post-hoc* analysis showed that D1-PYR in CUS mice exhibited significantly more depolarized RMP (−69.68 ± 1.75 mV) vs control mice (−65.60 ± 1.42 mV) (t_(16)_= −2.70, p=0.016), but no difference in D2-PYR (D2-PYR: CON −69.00 ± 2.78 mV; CUS −67.00 ± 1.30; t_(15)_= −0.624, p=0.54; data not shown). Previous research findings in L5 showing D1-PYR neurons are more hyperpolarized than D2-PYR (Seong & Carter, 2012), an inconsistency with our findings that may be due to the heterogeneity of pyramidal neurons within L5/6 of the mPFC (Kawaguchi, 1993; Yang et al., 1996). Furthermore, there was a significant condition (stress, naïve) by cell-type (D1-PYR, D2-PYR) interaction in PrL rheobase (F_(1,36)_= 12.38, p=0.001; Figure 3D). Examination of rheobase (Figure 3D) showed a significant interaction between experience and cell-type (F_(1,36)_=12.38, p=0.001). Post-hoc analysis showed that D1-PYR and D2-PYR rheobase was not significantly different in control mice, however CUS significantly reduced rheobase of D1-PYR compared to CON mice (CON: 95.00 ± 15.00 pA, CUS: 55.56 ± 5.56 pA; p=0.04). Alternatively, D2-PYR in CUS exposed mice exhibited a significantly higher rheobase compared to CON D2-PYR (CON: 83.64 ± 13.43 pA, CUS: 131.11 ± 12.52 pA, p=0.008). These changes indicate that the lack of effect on rheobase following CUS in randomly selected populations of cells likely reflects a bidirectional change in firing threshold among two distinct cell-types.

Examination of action potential frequency showed that although D1-PYR in CUS mice had a reduced firing threshold, frequency of firing was significantly reduced at higher current amplitude (interaction: F_(19,285)_= 14.56, p<0.001; Figure 3E). Conversely, comparison of firing in D2-PYR found only a main effect of condition, whereby CUS significantly reduced the overall firing frequency compared to cells from control mice (condition: F_(1,18)_= 4.36, p=0.05; interaction F_(19,342)_= 1.00, p=0.46; Figure 3F). These findings indicate that stress increases the likelihood of D1-PYR to fire, but reduces their firing capacity at more depolarized potentials, whereas CUS reduces activation of D2-PYR.

As no significant effects on firing were also observed in randomly selected IL PYR, we next examined whether stress produced a cell-specific change in firing properties based on D1- and D2R expression (Figure 3G). Here a combination of putative and fluorescently identified neurons was used, with no significant differences in the threshold to fire an action potential observed based on cell-type, condition, or a condition by cell-type interaction (condition: F_(1,38)_= 1.38, p=0.25; cell-type: F_(1,38)_=0.54, p=0.47; cell x condition: F_(1,38)_=0.03, p=0.87; Figure 3H). Additionally, no differences were observed in action potential frequency in either D1-PYR (condition: F_(1,12)_=4.04, p=0.07; condition x current F_(19,228)_=0.79, p=0.71; Figure 3I) or D2-PYR (condition: F_(1, 24)_= 0.10, p=0.75; condition x current (F_(19, 456)_= 1.11, p=0.33; Figure 3I).

### 3.4 Cell-specific effects of CUS on PrL D1- and D2-PYR synaptic transmission

Functional integrity of mPFC information processing relies on a dynamic balance of excitatory and inhibitory transmission (Isaacson & Scanziani, 2011; Kim et al., 2016; Sohal et al., 2009; Yizhar et al., 2011). Recent evidence indicates that sub-types of mPFC PYR receive distinct forms of excitatory and inhibitory regulation (Lee et al., 2014). To further explore CUS dependent cell-specific plasticity, we performed *ex vivo* recordings to examine changes in miniature excitatory (mEPSC) and inhibitory (mIPSC) currents in the PrL – a direct and selective measure of synaptic AMPA and GABA_A_ receptor function, respectively (Figure 4). In the current study, we found no significant differences in the amplitude of mEPSCs (Figure 4A-C; D1-PYR: 10.73 ± 0.83pA, D2-PYR: 10.97 ± 0.50 pA) or mIPSCs (Figure 4E-H; D1-PYR: 14.13 ± 0.84 pA, D2-PYR: 13.84 ± 1.14 pA) in controls. CUS did not alter the amplitude of mEPSCs (Figure 4C) or mIPSCs (Figure 4G) based on a lack of condition, cell-type, or cell-type x condition interaction (condition: F_(1,37)_=0.008, p=0.93; cell-type: F_(1,37)_=1.02, p=0.32; cell x condition: F_(1,37)_=1.67, p=0.21).

**Figure 4.**
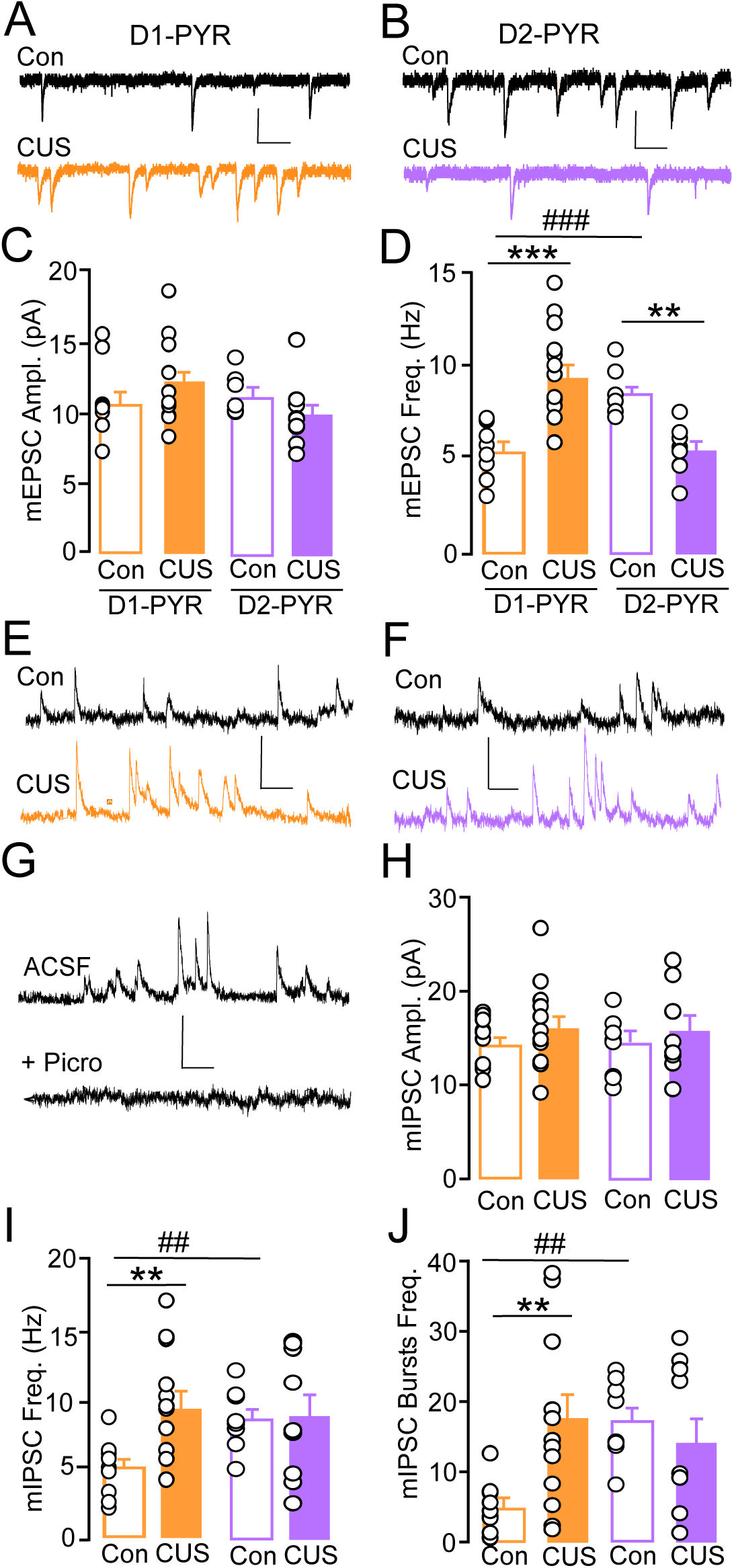
(A, B) Representative traces of mEPSC from CON and CUS mice. (A) D1-PYR (black, orange) and (B) D2-PYR (black, purple) in the PrL. (C) There were no differences in mean mEPSC amplitude across cell-type or condition. (D) In CON mice, mean frequency of mEPSCs was significantly higher in D2-PYR compared to D1-PYR. CUS increased mean mEPSC frequency in D1-PYR and reduced it in D2-PYR (*n*=10-12/group, *N*=6-7/group). (E, F) Representative traces of mIPSC from CON and CUS mice. (E) D1-PYR and (F) D2-PYR in the PrL. (G) Representative mIPSC traces under ACSF alone versus lack of events following subsequent application of the selective GABA_A_ antagonist, picrotoxin (Picro). (H) There were no significant differences in mean mIPSC amplitude across cell-type or condition. (I) Mean mIPSC frequency and (J) mean bursts of mIPSCs (4+events/150 ms) per sweep was significantly higher in D2-PYR compared to D1-PYR in CON mice. Mean mIPSC event and burst frequency was significantly greater in D1-PYR of CUS vs CON mice, while no difference was observed in D2-PYR (*n=*10-12/group, *N*=7/group). mEPSC scale bar, 20pA/100msec; mIPSC scale bar, 30 pA/100. **p<0.01, ***p<0.001 CUS versus CON; ^##^ p<0.01, ^###^ p<0.001 D1-PYR CON versus D2-PYR CON.

Alternatively, we found a significant interaction of cell-type and condition on mEPSC frequency (F_(1,37)_=32.31, p<0.001). D2-PYR from control mice showed significantly higher mEPSC frequency compared to D1-PYR (CON D1-PYR: 5.26 ± 0.47 Hz, CON D2-PYR: 8.28 ± 0.43 Hz, p<0.001; Figure 4D). Compared to controls, CUS significantly increased mEPSC frequency at D1-PYR (CUS 9.14 ± 0.72 Hz, p<0.001), while reduced frequency at D2-PYR (CUS 5.44 ± 0.59 Hz; p=0.003; Figure 4D). A similar interaction of condition and cell-type was observed for mIPSC frequency (F_(1,38)_=5.70, p=0.023) and frequency of IPSC bursts per sweep (F_(1,38)_=8.51, p=0.006). *Post hoc* analysis showed that under control conditions, D2-PYR exhibit higher mIPSC frequency (CON D1-PYR: 4.78 ± 0.61 Hz, CON D2-PYR: 9.39 ± 0.77 Hz; p<0.01) and burst frequency (CON D1-PYR: 4.24 ± 1.17, CON D2-PYR: 17.48 ± 2.01; p<0.004) compared to D1-PYR (Figure 4I, J). Compared to controls, CUS significantly increased mIPSC frequency and burst frequency in D1-PYR (frequency: 9.14 ± 0.72 Hz; burst frequency: 16.83 ± 2.01 bursts; Figure 4J). CUS did not alter frequency or burst frequency in D2-PYR (frequency: 8.08 ± 1.21; burst frequency: 12.31 ± 3.32 bursts). Taken together, these data show that D1- and D2-PYR maintain distinctly different excitatory and inhibitory synaptic regulation under naïve conditions, and that CUS promotes opposing changes in excitatory and inhibitory drive at D1- and D2-PYR that may contribute to divergent effects on intrinsic excitability, and that these effects may result in secondary adaptations in inhibitory signaling at D1-PYR.

## 4.0 Discussion

The current study demonstrated that using a chronic unpredictable model of stress with increased variability and enhanced intensity of stressors produced anhedonia-anxiety-like behavior and increased passive coping strategy in a significantly shorter time frame (2-4 weeks) than previous reports using a single daily exposure to CUS in C57BL6 mice (Monteiro et al., 2015). We find that under control conditions, PrL D1- and D2-PYR did not exhibit distinctions in firing threshold, but showed distinctions in excitatiory:inhibitory synaptic drive, and that our model of CUS exposure produced enduring and opposing adaptations in both intrinsic physiology and synaptic regulation of these pyramidal subpopulations. D1-PYR from CUS mice exhibited a reduction in firing threshold (increased excitability) but impaired maintenance of firing capacity at more depolarized potentials that was paralleled by enhanced frequency of excitatory AMPAR-mEPSCs and inhibitory GABA_A_-mEPSCs. Alternatively, CUS promoted an increase in D2-PYR firing threshold (reduced excitation) that was paralleled by reductions in excitatory drive. Taken together, these results build upon previously identified intrinsic differences in D1- and D2-PYR and demonstrate for the first time, that prolonged stress may produce abnormalities in PFC-dependent behavior by uniquely modifying mPFC circuits comprised of D1- and D2-PYR, and that these modifications reflect overlapping and distinct forms of plasticity.

### 4.1 Impact of stress intensity and predictability on affective behavior

The influence of chronic stress exposure on affective behaviors has been well-established in rats, with a variety of mild, unpredictable, and social stress paradigms able to reduce sucrose preference, as well as increase anxiety- and depression-like behavior (Vasconcelos, Stein, & de Almeida, 2015; Willner, 2017). Alternatively, inherent strain differences in stress susceptibility have presented a challenge towards the use of mice – an approach that would greatly expand genetic manipulations and cell-specific identification through the use of transgenic animals. Recent work by Monteiro and colleagues (2015) laid the groundwork for establishing mouse-specific CUS protocols that produce reliable deficits in affective behavior, however these protocols involved up to eight weeks of once daily stress exposure – a length of time that not only reduces throughput, but negates advantages related to per diem costs associated with housing mice versus rats. As unpredictable stressors often more negatively impact humans compared to predictable ones (Anisman & Matheson, 2005; Bale, 2005; Kendler et al., 1998, 1999; Moghaddam & Javitt, 2012; Willner & Mitchell, 2002), it was plausible that manipulations of intensity and predictability would allow for the reduction in length of stress exposure, while resulting in similar behavioral phenotypes. Similar to CUS models in rats, the current protocol utilized two daily stressors, but sought to increase overall stress exposure intensity by combining stressors, using stressors with greater intensity (e.g. forced swim and restraint) more frequently, and decreasing predictability by using multiple distinctly different contexts and varying the time of day in which stressors were given. These changes reduced time spent in the open arm of an EPM, reduced preference for sucrose, and increased passive coping strategies – the latter of which persisted around 17 days following stress exposure – as indicative of increased anxiety-, anhedonia- and depression-like behavior. Notably, our unpublished data indicate that reductions in open arm time following 2 weeks of more intense stress were no longer present at 17-21 days post-stress, suggesting that our CUS protocol produces enduring deficits in depression-but not anxiety-like behaviors. Our data support the ability to use C57BL6 mouse models to study plasticity associated with CUS without drastically prolonging stress exposure, however, as previous reports have shown a delayed emergence of affect behavior following chronic stress exposure in rats (Matuszewich et al., 2007), it will be important for future studies to characterize the timeline of the behavioral and physiological changes produced by this CUS regimen

### 4.2 Bidirectional changes in prelimbic D1- and D2-PYR intrinsic excitability

Neuroanatomical studies indicate that similar to medium spiny neurons in the striatum, mPFC pyramidal neurons can be canonically divided based on the expression of D1 or D2 receptors (Gaspar et al., 1995; Santana et al., 2009; Vincent et al., 1993). Pharmacological evidence indicates that mPFC dopamine D1 and D2 receptors modulate dissociable (often opposing) aspects of cognitive and affective behavior (Bai et al., 2017; Jenni et al., 2017; Sawaguchi & Goldman-Rakic, 1994; Seamans, Floresco, & Phillips, 1998; Shinohara et al., 2018), and distinct PFC circuits (Durstewitz, Seamans, & Sejnowski, 2000; Durstewitz, Vittoz, Floresco, & Seamans, 2010; Jenni et al., 2017; Seamans & Yang, 2004). More recent evidence utilizing optogenetics, chemogenetics, and D1- or D2-cre transgenic mouse lines supports this notion, with numerous studies demonstrating that manipulating activity of PFC D1 and D2R-expressing cell-bodies produces distinct modifications in behavior related to feeding, social interaction, and depression-associated behavior, that may not be reproducible with general population manipulations (Brumback et al., 2018; Hare et al., 2019; Land et al., 2014; Narayanan, Land, Solder, Deisseroth, & DiLeone, 2012; Shinohara et al., 2018). A number of these studies have directly demonstrated that D1-PYR networks are specifically involved in regulating this behavior through downstream terminal stimulation approaches (Hare et al., 2019; Land et al., 2014) however as D1- and D2R have been found pre- and postsynaptically, as well as on principle and interneuron populations (Anastasiades et al., 2018; Benes et al., 1993; Santana et al., 2009; Vincent et al., 1993) and recent findings indicate that stimulation of D1- *and* D2R on PYR exert an excitatory effect (Gee et al., 2012; Robinson & Sohal, 2017; Seong & Carter, 2012), approaches involving intra-cranial pharmacological manipulations as well as intra-PFC manipulations in Cre-mice should be interpreted cautiously.

Studies of patients and animal models suggest that a functional imbalance in the ratio of PFC cellular excitation:inhibition causally underlies impaired working memory, social withdrawal, and anxiety-like behavior in stress-related psychiatric disorders (Fuchs et al., 2016; Gandal et al., 2012; Gonzalez-Burgos & Lewis, 2012; Holmes & Wellman, 2009; Javitt et al., 2011; Matsuo et al., 2007; Moghaddam & Javitt, 2012; Sohal et al., 2009; Yizhar et al., 2011). Intrinsic membrane properties play a critical part in determining this balance, as they directly shape neuronal output by influencing the probability of a neuron firing an action potential in response to excitatory synaptic inputs and modulate firing capacity. The lack of significant effect of stress on rheobase and action potential firing frequency in randomly selected populations in the current study closely resembles findings in adult male rats 24 h following conclusion of social defeat stress (Urban and Valentino, 2017). Thus, it is possible that null effects following social stress also reflect an opposing reduction and increase in firing thresholds (rheobase) in D1- and D2-PYR, respectively.

The underlying intrinsic mechanisms contributing to alterations in firing threshold remain unclear. Our previous work has shown that G protein-gated inwardly rectifying K+ channels (Girks) mediate most of the GABA_B_R-dependent inhibition of Layer 5/6 PYR in the mPFC, essentially acting as a neuronal off switch (M. Hearing et al., 2013; M. C. Hearing, Zink, & Wickman, 2012). Given that knockout of these channels or experience-dependent suppression of this signaling reduces firing thresholds akin to that observed in D1-PYR here, it is possible that CUS promotes a downregulation of GABA_B_R-Girk signaling in D1-PYR and perhaps an upregulation in D2-PYR. In addition to reduced firing threshold in D1-PYR, CUS promotes an apparent depolarization-induced blockade at higher current injections. As activity-dependent Girk channel plasticity has recently been implicated in the transition between tonic and burst firing modes in midbrain dopamine neurons, it is also possible that alterations in Girk channels could also contribute to the stress-related reduction in firing capacity (Lalive et al., 2014). Alternatively, this may reflect impaired hyperpolarization-activated graded persistent activity responsible for maintaining a constant firing rate or increased inhibitory transmission, as elevations in the frequency and bursting of mIPSCs were observed in D1-PYR (Winograd, Destexhe, & Sanchez-Vives, 2008). Regardless, as persistent activity in PFC networks is thought to be important for memory formation, conditioned associations, and working memory (Gilmartin & Helmstetter, 2010; Gilmartin, Kwapis, & Helmstetter, 2012; Gilmartin & McEchron, 2005; Gilmartin, Miyawaki, Helmstetter, & Diba, 2013; Kwapis, Jarome, & Helmstetter, 2014; Runyan, Moore, & Dash, 2004; Seamans, Nogueira, & Lavin, 2003) it is possible that deficits in these facets of cognition may be due in part to reduced firing capacity in D1- or D2-PYR.

### 4.3 Effects of CUS on infralimbic pyramidal neuron excitability

Although the present study identified adaptations in D1- and D2-PYR within the PrL that were not initially observed when examining a general population of cells, data obtained from putative and fluorescently identified D1- and D2-PYR in the IL did not show significant differences in intrinsic excitability. The lack of effect on IL intrinsic excitability is particularly surprising given previous work demonstrating that CUS increases inhibitory input and transmission, that is paralleled by increased inhibitory appositions on glutamatergic neurons when sampling from unidentified populations of IL PYR (McKlveen et al., 2016).

Although outside the scope of the current study, it is possible that stress promotes modifications in the IL that are not only cell-specific but also pathway specific, as stress is known to alter basolateral amygdala (BLA)-to-PFC input without influencing BLA inputs projecting to the bed nucleus of the stria terminalis (BNST) (Lowery-Gionta et al., 2018). Circuit specific structural modifications have also been implicated in apical dendrite retraction following chronic restraint stress in IL circuits, as a population of Layer 2/3 PYR projecting to the BLA appear to be spared from stress plasticity (Shansky & Morrison, 2009). These findings indicate that stress almost assuredly promotes enduring plasticity in IL neurons, however these modifications may be more complex and/or specific based on anatomical connectivity – a possibility we are currently exploring.

### 4.4 Cell-specific effects of stress on synaptic transmission

Although stress-induced structural plasticity in the mPFC has been recognized for more than a decade as a prominent factor in PFC dysfunction few studies have examined the pathway- and cell-specific locus of these adaptations (Lowery-Gionta et al., 2018; McEwen & Morrison, 2013; McKlveen et al., 2016; Shansky & Morrison, 2009; Yuen et al., 2012). Previous work has shown that in early adolescent male mice, CUS transiently reduces PrL PYR glutamate receptor expression and excitatory synaptic transmission which returned to control levels by day 5 post stress, although the exact population of pyramidal neurons (layer 2/3 or layer 5/6) examined was not apparent (Yuen et al., 2012). It is possible that the apparent lack of enduring change in excitatory plasticity reflects examination of unidentified populations, as we observed divergent effects on mEPSC frequency in D1 vs D2-PYR. Alternatively, early reductions in glutamate signaling may represents a generalized permissive functional plasticity that precedes divergent adaptations (Kourrich, Calu, & Bonci, 2015). It is also possible that early reductions in excitatory signaling is specific for adolescents, as PrL PYR from mid-adolescent but not adult male rats, exhibit reductions in excitatory transmission 24 h following prolonged social defeat stress (Urban and Valentino, 2017). The current findings of increased mEPSC frequency but not amplitude in D1-PYR tangentially aligns with other findings of increased presynaptic glutamate release in BLA to PFC synapses, with no differences in AMPA/NMDA ratio noted (Lowery-Gionta et al., 2018). Given the opposing effects of CUS on the frequency of mEPSCs in D1-PYR but not D2-PYR, it is possible that social defeat and restraint did alter excitatory transmission in adults, albeit in opposing fashion.

Recent work has shown that type A and B PYR in the PrL exhibit properties akin to D2- and D1-PYR, respectively, and that these populations receive distinct forms of input that may sub-serve divergent functions (Gee et al., 2012; Lee et al., 2014; Seong & Carter, 2012). For example, type A PYR (i.e., D2-PYR) are preferentially inhibited by fast-spiking parvalbumin internueurons but not somatostatin interneurons (Lee et al., 2014). Our findings build on these observations, showing elevated excitatory and inhibitory signaling in D2-vs D1-PYR – highlighting a need to understand how these intrinsic differences contribute to CUS-dependent plasticity will be an important future direction. Interestingly, we find that increases in excitatory transmission in D1-PYR were paralleled by enhanced inhibitory drive. Although the source of inhibitory change is unclear, local neocortical circuitry includes collateral connections between pyramidal neurons as well as reciprocal connections between pyramidal and local inter-neurons (Isaacson & Scanziani, 2011), thus it is possible that increased excitation of D1-PYR increases activation of and transmitter release from local GABA neurons – a possibility that aligns with the selective increase in mIPSC frequency and bursting rather than amplitude (Sohal, 2012). As all of our observed synaptic adaptations were selective for changes in the frequency of PSCs, it will also be important to determine whether these reflect presynaptic or postsynaptic structural modifications.

### 4.4 Functional Implications

Emerging evidence indicates that PFC-dependent regulation of cognition and affective behavior are governed through a complex array of cortical networks comprised of neuronal subpopulations that express innate physiological and synaptic properties that likely determine how they influence behavior and undergo plasticity (McEwen & Morrison, 2013). It is tempting to speculate that the divergent effects on D1- and D2-PYR in the current study underlie specific stress-related pathologies. For example, reductions in the output of D2-PYR may reflect reduced top-down control over affect related behavior, and thus permit the emergence of increased anxiety- and depression-like behavior. On the other hand, recent findings indicate that cognitive deficits in a number of disorders may reflect a shift towards cortical excitation and increased (disorganized) firing of mPFC PYR that disrupts cortical information flow and cognitive performance (Fuchs et al., 2016; Gonzalez-Burgos & Lewis, 2012; Javitt et al., 2011; Matsuo et al., 2007; Moghaddam & Javitt, 2012).

## 5.0 Conclusion

The ability of ostensibly unrelated disorders to give rise to seemingly similar psychiatric phenotypes highlights a need to identify circuit-level concepts that could unify diverse factors under a common pathophysiology. The current work indicates that stress-related pathophysiology likely manifests through dynamic alterations in communication within specific cortical networks. Our findings highlight the need to gain a better understanding of which neural pathways are responsible for regulating behavior under naïve and pathological states to identify more targeted approaches to effectively treat neuropsychiatric disorders that encompass varied, co-occurring symptoms, and divergent responses to treatment.

## Acknowledgements

These studies were supported by funding from the Brain and Behavior Research Foundation (#26299), Marquette University Regular Research Grant, and the Charles E Kubly Mental Health Research Foundation at Marquette University.

